# Ketamine disrupts neuromodulatory control of glutamatergic synaptic transmission

**DOI:** 10.1101/341172

**Authors:** Gyorgy Lur, Michael J. Higley

## Abstract

A growing body of literature has demonstrated the potential for ketamine in the treatment of major depression. Sub-anesthetic doses produce rapid and sustained changes in depressive behavior, both in patients and rodent models, associated with reorganization of glutamatergic synapses in the prefrontal cortex (PFC). While ketamine is known to regulate NMDA-type glutamate receptors (NMDARs), the full complement of downstream cellular consequences for ketamine administration are not well understood. Here, we combine electrophysiology with 2-photon imaging and glutamate uncaging in acute slices of mouse PFC to further examine how ketamine alters glutamatergic synaptic transmission. We find that four hours after ketamine treatment, glutamatergic synapses themselves are not significantly affected. However, expression levels of the neuromodulatory Regulator of G-protein Signaling (RGS4) are dramatically reduced. This loss of RGS4 activity disrupts the normal compartmentalization of synaptic neuromodulation. Thus, under control conditions, α2 adrenergic receptors and GABA_B_ receptors selectively inhibit AMPA-type glutamate receptors (AMPARs) and NMDARs, respectively. After ketamine-induced loss of RGS4 activity, this selectivity is lost, with both modulatory systems broadly inhibiting glutamatergic transmission. These results demonstrate a novel mechanism by which ketamine can influence synaptic signaling and provide new avenues for the exploration of therapeutics directed at treating neuropsychiatric disorders, such as depression.

## Introduction

Major depression, with a lifetime prevalence of 17%, presents a significant psychological and economical burden for both individuals and society (Kessler *et al*, 2003; Kessler *et al*, 2005). Despite enormous efforts to develop effective treatments, available therapeutic interventions have considerable limitations. For example, most antidepressant medications take several weeks to achieve maximal benefit and a significant fraction of patients remain refractory to treatment (Fava *et al*, 2008; Trivedi *et al*, 2006). However, recent studies in both clinical and basic science fields have demonstrated promising results using low doses of the drug ketamine. Indeed, sub-anesthetic doses of ketamine produce rapid antidepressant actions within a few hours (Berman *et al*, 2000; Krystal *et al*, 2013; Zarate *et al*, 2006), even in otherwise refractory patients (Mathew *et al*, 2012). Thus, the potential benefits of this new pharmacological intervention provide great promise for the treatment of major depression.

Surprisingly, the neurological mechanisms underlying the antidepressant actions of ketamine remain poorly understood. Recent efforts have focused on the regulation of glutamatergic synapses in the prefrontal cortex (PFC) as a potential process by which ketamine modulates behavior. Ketamine itself is an antagonist of NMDA-type glutamate receptors (NMDARs)(MacDonald *et al*, 1987), though it may have other actions as well (Sleigh *et al*, 2014). Acute administration of ketamine produces mild dissociative effects that subside within two hours after administration (Krystal *et al*, 1994), while the antidepressant actions may persist for up to a week (Berman *et al*, 2000; Zarate *et al*, 2006). In animal models, ketamine stimulates a signaling cascade that produces long-term enhancement of glutamatergic transmission in the PFC, including increased synaptic protein synthesis and increased density of dendritic spines, the structural sites of individual excitatory inputs (Duman and Aghajanian, 2012; Holtmaat and Svoboda, 2009; Li *et al*, 2010; Moghaddam *et al*, 1997). Moreover, these synaptic changes persist for several days after administration (Li *et al*, 2010).

The cellular mechanisms underlying these alterations in synaptic structure and function are unclear, and a variety of signaling pathways have been implicated in linking NMDAR blockade to long-term alteration of glutamatergic signaling, including the mammalian target of rapamycin (mTOR) and brain-derived neurotrophic factor (BDNF) (Autry *et al*, 2011; Li *et al*, 2010; Liu *et al*, 2012). Both these processes have been shown to regulate the growth and stability of glutamatergic synapses (Jourdi *et al*, 2009; Takei *et al*, 2004). However, ketamine has also been linked to alterations in inhibitory GABAergic signaling, both directly and indirectly by disrupting activity in inhibitory interneurons (Homayoun and Moghaddam, 2007; Kavalali and Monteggia, 2012; Wohleb *et al*, 2017).

A recent study suggested that acute administration of ketamine might also impact the function of the protein regulator of G-protein signaling type-4 (RGS4) (Stratinaki *et al*, 2013). RGS4 is a GTPase activating protein that accelerates the hydrolysis of GTP to GDP following the activation of G proteins by a variety of ligand-receptor interactions (Bansal *et al*, 2007; Watson *et al*, 1996). Behavioral studies in mice lacking RGS4 demonstrated that this enzyme can act as a key negative modulator of ketamine-mediated antidepressant actions (Stratinaki *et al*, 2013), and ketamine itself is capable of reducing expression levels of RGS4 in the PFC. We previously found that RGS4 plays a crucial role in regulating the neuromodulation of glutamatergic synapses in the PFC (Lur and Higley, 2015). Our data showed that the specificity of adrenergic and GABAergic control over AMPA- and NMDA-type glutamate receptors, respectively, is abolished following pharmacological blockade of RGS4 function.

As neuromodulation may play a key role in normal prefrontal function, we investigated whether ketamine might disrupt the regulation of glutamate receptors in the PFC. Using a combination of electrophysiology, 2-photon imaging, and glutamate uncaging in acute slices from the mouse PFC, we found that a single dose of ketamine did not alter basal function of postsynaptic glutamate receptors. However, ketamine did produce a substantial reduction in prefrontal RGS4 expression in the PFC and abolished the specificity of adrenergic and GABAergic modulation of excitatory synapses. Thus, our work suggests a novel mechanism by which acute ketamine can influence glutamatergic signaling and potentially contribute to its antidepressant actions.

## Materials and Methods

### Animals and drug treatment

All animal handling was performed in accordance with guidelines approved by the Yale Institutional Animal Care and Use Committee and federal guidelines. Wild-type C57/Bl6 mice (P22-36) were injected i.p. with either saline vehicle or ketamine (15 mg/kg) 4 hours prior to experiments.

### Slice Preparation

For glutamate uncaging experiments, we prepared acute prefrontal cortical (PFC) slices as previously described (Lur *et al*, 2015). Briefly, mice were anesthetized with isoflurane and decapitated, and coronal slices (300 μm) were cut in ice-cold solution containing (in mM): 110 choline, 25 NaHCO_3_, 1.25 NaH_2_PO_4_, 3 KCl, 7 MgCl_2_, 0.5 CaCl_2_, 10 glucose, 11.6 sodium ascorbate and 3.1 sodium pyruvate, bubbled with 95% O_2_ and 5% CO_2_. Slices containing the prelimbic-infralimbic regions of the PFC were then transferred to artificial cerebrospinal fluid (ACSF) containing (in mM): 126 NaCl, 26 NaHCO_3_, 1.25 NaH_2_PO_4_, 3 KCl, 1 MgCl_2_, 2 CaCl_2_, 10 glucose, 0.4 sodium ascorbate, 2 sodium pyruvate and 3 myo-inositol, bubbled with 95% O_2_ and 5% CO_2_. After an incubation period of 15 min at 34°C, the slices were maintained at 22–24°C for at least 20 min before use.

### Electrophysiology and imaging

All experiments were conducted at near physiological temperature (32-34°C) in a submersion-type recording chamber. Whole-cell recordings in voltage clamp mode were obtained from layer 5 pyramidal cells (400- 500 μm from the pial surface) identified with infrared differential interference contrast. Glass electrodes (1.8-3.0 MΩ) were filled with internal solution containing (in mM): 135 CsMeSO_3_, 10 HEPES, 4 MgCl_2_, 4 Na_2_ATP, 0.4 NaGTP, 10 sodium creatine phosphate and 0.2% Neurobiotin (Vector Laboratories) adjusted to pH 7.3 with CsOH. Red-fluorescent Alexa Fluor-594 (10 μM, Invitrogen) and the green-fluorescent calcium (Ca2+)-sensitive Fluo-5F (300 μM, Invitrogen) were included in the pipette solution. Neurons were filled via the patch electrode for 10 min before imaging. Series resistance was 10-22 MΩ and uncompensated. Electrophysiological recordings were made using a Multiclamp 700B amplifier (Molecular Devices), filtered at 4 kHz, and digitized at 10 kHz. Typically, 7-10 trials were averaged into each response.

2-photon imaging was accomplished with a custom-modified Olympus BX51-WI microscope (Olympus, Japan), including components manufactured by Mike’s Machine Company (Higley and Sabatini, 2010). Fluorophores were excited using 840 nm light from a pulsed titanium-sapphire laser (Ultra2, Coherent). Emitted green and red photons were separated with appropriate optics (Chroma, Semrock) and collected by photomultiplier tubes (Hamamatsu).

For Ca2+ imaging, signals were collected during 500 Hz line scans across a spine. Ca2+ signals were first quantified as increases in green fluorescence from baseline normalized to the average red fluorescence (AG/R). We then expressed fluorescence changes as the fraction of the G/R ratio measured in saturating Ca2+ (ΔG/G_sat_)(Lur and Higley 2015).

### 2-Photon Glutamate uncaging

For focal stimulation of single dendritic spines, we used 2-photon laser uncaging of glutamate (2PLU). To photorelease glutamate, a second Ti-Sapphire laser tuned to 720 nm was introduced into the light path using polarization optics. Laser power was calibrated for each spine by directing the uncaging spot to the middle of the spine head. We adjusted uncaging power to achieve 50% photobleaching of the Alexa 594 dye filling the spine (Lur *et al*, 2015). The power used for 2PLU ranged from 8 to 25 mW, pulse duration was 2 ms. For synaptic stimulation, we typically uncaged glutamate at 3-4 separate locations around a single spine head to find a “hot spot”, the place of the largest response. Our previous measurements indicate that this stimulus resulted in uEPSCs similar in size and kinetics to spontaneous mEPSCs (Lur *et al*, 2015).

### Data acquisition and analysis

Imaging and physiology data were acquired using National Instruments data acquisition boards and custom software written in MATLAB (Mathworks, (Pologruto *et al*, 2003)). Off-line analysis was performed using custom routines written in MATLAB and IgorPro (Wavemetrics). AMPAR-mediated EPSC amplitudes were calculated by finding the peak of the current traces and averaging the values within a 0.3 ms window. NMDAR-mediated currents were measured in a 3 ms window around the peak. 2PLU-evoked ΔCa2+ was calculated as the average ΔG/G_sat_ over a 100 ms window, starting 5 ms after the uncaging event. Statistical comparisons were conducted in GraphPad Prism 5. All data were analyzed using one-way analysis of variance (ANOVA)-tests corrected for multiple comparisons (Tukey).

### Pharmacology and reagents

2PLU experiments were performed in normal ACSF supplemented with MNI-glutamate (2.5 mM) and D-serine (10 μM). To isolate AMPAR-mediated currents in voltage clamp experiments, we added TTX (1 μM) to block sodium channels, picrotoxin (50 μM) to block GABA_A_ receptors, CGP55845 (3 μM) to block GABA_B_ receptors, and CPP (10 μM) to block NMDA-type glutamate receptors to the ACSF. To isolate NMDAR-mediated currents, we modified our original ACSF to contain 0 mM Mg and 3 mM Ca2+ and included TTX (1 μM), picrotoxin (50 μM), CGP55845 (3 μM), and NBQX (10 μM) to block AMPA-type glutamate receptors. GPCR agonists were applied 5-7 minutes prior to data collection. All compounds and salts were from Tocris and Sigma, respectively.

### Western blot analysis

For RGS4 western blot analysis, we prepared 300 μm thick brain slices containing the PFC from p22-42 C57/bl6 mice as described above. Following the recovery period, the prelimbic region of the prefrontal cortex was dissected out of the slices on ice. Tissue samples were homogenized and sonicated in ice cold lysis buffer containing 20 mM Tris, 1 mM EDTA and 1x Halt protease and phosphatase inhibitor cocktail (Thermo Scientific) and 0.5% SDS, pH 8.0. After a 10-minute centrifugation at 14000 rpm, the supernatant was collected, and protein content was determined using Pierce BCA Protein Assay (Thermo Scientific). Samples containing equal amounts of protein were separated on a 6% poly-acrylamide gel and transferred to PVDF membranes. After blocking for 1h at room temperature with 3% non-fat milk and 0.02% Na-azide in Tris buffered salt solution with 0.05% Tween 20 (TBST), membranes were immunoreacted with primary antibody against RGS4 (Millipore, RBT17) in 1% milk and 0.02% Naazide in TBST, 1:1000, overnight. After washing off excess primary antibody and incubation with the appropriate HRP conjugated secondary antibody (GE Healthcare, UK) for 2 hours at room temperature in TBST, bands were visualized using HyGlo Chemiluminescent HRP Antibody Detection Reagent (Denville Scientific Inc.) and exposed onto autoradiography film (Denville Scientific Inc.). Membranes were then stripped from antibodies using Restore Plus Western Blot Stripping Buffer (15 minutes at room temperature, Thermo Scientific), re-blocked and immunoreacted with anti-β-tubulin (SIGMA) primary antibody followed by the appropriate HRP-secondary antibody to establish total amount β-tubulin in the samples. Autoradiography films were developed in a Kodak automatic developer, then scanned and analyzed with ImageJ. RGS4 expression was quantified as RGS4 / β-tubulin.

## Results

To investigate the consequences of ketamine for glutamatergic signaling, we injected mice with either a single dose of ketamine (15 mg/kg) or saline vehicle. At four hours post-injection, we prepared acute brain slices from the medial prefrontal cortex (PFC) of the injected animals and made whole cell voltage clamp recordings from layer 5 pyramidal neurons in the prelimbic region (Figure 1A). We measured excitatory postsynaptic currents (EPSCs) evoked by 2-photon laser uncaging (2PLU) of glutamate onto spines along the basal dendrites, while simultaneously monitoring intra-spine Ca2+ transients using 2-photon laser scanning microscopy (2PLSM)(Figure 1B). We found no difference in AMPAR-mediated EPSCs between vehicle (20.9 ± 1.9 pA, n= 31 spines) and ketamine (21.0 ± 1.4 pA, n=33 spines, p=0.96, t-test, Figures 1C and 4A) groups. Notably, we also failed to observe alterations in NMDAR-mediated currents (vehicle: 22.9 ± 2 pA, n=31 spines, ketamine: 21.4 ± 1.8 pA, n=31 spines, p=0.59, t-test) or Ca2+ transients (vehicle: 0.52 ± 0.02 ΔG/Gsat, ketamine: 0.54 ± 0.02 ΔG/G_sat_, p=0.31, t-test, Figures 1C and 4B), indicating the absence of persistent NMDAR blockade. Overall, our results suggest that ketamine produces minimal postsynaptic change in glutamatergic synapses at this time point.

**Figure 1.**
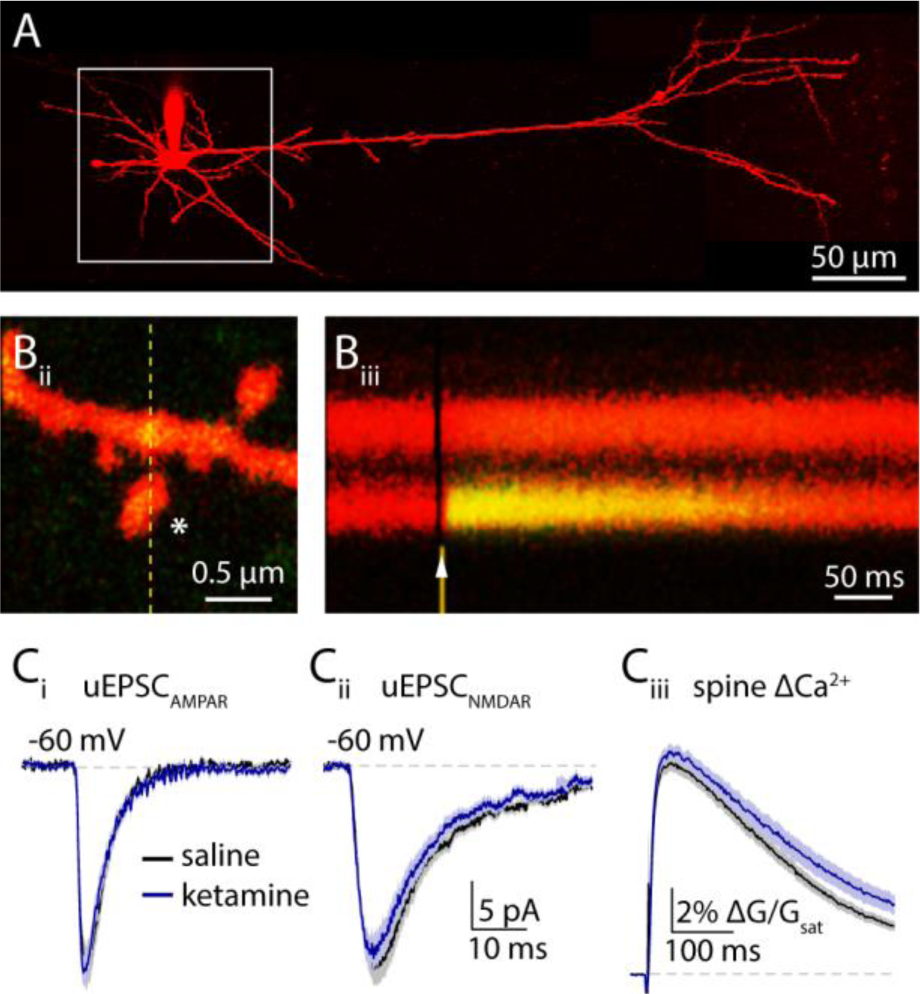
Ketamine does not affect glutamatergic transmission 4 hours post treatment. (A) A 2-photon image of a layer 5 pyramidal neuron visualized by Alexa 594 flourescence. White box highlights the extent of the basal dendritic arbor searched for spines. (Bi) Image of a dendritic spine. White asterisk shows the uncaging location. Fluorescence intensity was measured in a line scan highlighted by the yellow dashed line. (Bii) Chronogram showing the change in green fluorescence indicating a transient increase of intracellular Ca2+ concentration in the spine head in response to glutamate uncaging (arrowhead). (Ci) Mean AMPAR-mediated uEPSCs in vehicle (black) and in ketamine (blue) treated animals ± SEM (shaded areas). (Cii) 2PLU-evoked NMDAR currents and (Ciii) Ca2+ transients in vehicle (black) and in ketamine treated animals (blue), mean (solid lines) ± SEM (shaded areas). *: p<0.05, unpaired t-test.

Endogenous control of glutamatergic transmission by GPCRs plays a central role in prefrontal function. We previously showed that both noradrenergic and GABAergic modulation of glutamatergic synapses is influenced by the activity of the small GTPase RGS4 (Lur *et al*, 2015), whose expression levels are thought to be regulated by antidepressants including ketamine (Stratinaki *et al*, 2013). To examine the consequences of acute ketamine on RGS4 function in the prefrontal cortex, we first prepared tissue samples from the prelimbic region four hours after animals were injected with either ketamine or saline. Western-blot analysis confirmed a 50 ± 0.1% reduction of RGS4 protein expression (p=0.0016, n=5 animals, unpaired t-test, Figure 2A and B), suggesting that ketamine may disrupt neuromodulation at glutamatergic synapses.

**Figure 2.**
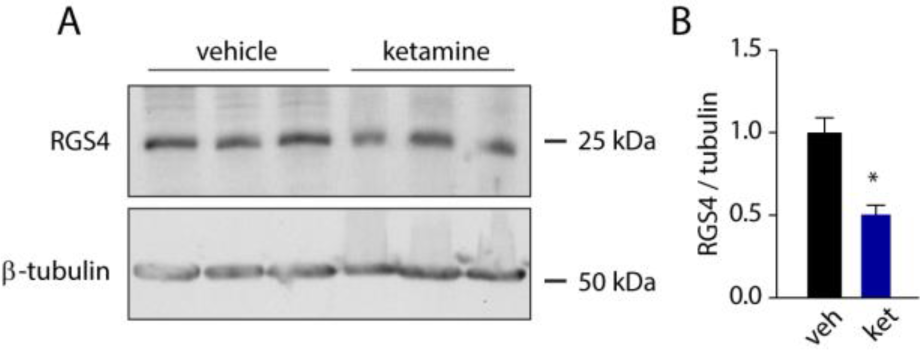
Acute ketamine treatment reduces prefrontal RGS4 expression. (A) Example western blot of RGS4 and β-tubulin in vehicle- versus ketamine-treated mice. (B) Quantification of RGS4 levels in vehicle (black) and ketamine (blue) treated mice. Bars show mean ± SEM, *: p<0.05, unpaired t-test.

Activation of alpha2 adrenergic receptors and GABAB receptors negatively modulate AMPARs and NMDARs, respectively via downregulation of protein kinase A (PKA) activity (Lur *et al*, 2015). Moreover, this selective coupling of GPCRs to distinct synaptic proteins requires RGS4 and is lost following small molecule antagonism of RGS4 activity (Lur *et al*, 2015). We therefore asked whether ketamine produces similar dysregulation of synaptic modulation. First, we confirmed that, in saline-injected mice, application of the alpha2 adrenergic agonist guanfacine significantly reduced 2PLU-evoked AMPAR-mediated currents (to 13.0 ± 0.8 pA, n=25 spines, one-way ANOVA (F=9.93, p<0.0001), Tukey’s multiple comparison test p<0.01) but did not alter 2PLU-evoked NMDAR-mediated currents (22.3 ± 2 pA, n=33 spines, one-way ANOVA (F=6.53, p=0.0004), Tukey’s multiple comparison test p>0.05, Figures 3Aii and 4B) or ΔCa2+ (0.54 ± 0.02 ΔG/G_sat_, one-way ANOVA (F=8.1, p<0.0001), Tukey’s multiple comparison test p>0.05, Figures 3A, 4). Conversely, application of the GABAB agonist baclofen did not alter 2PLU-evoked AMPAR-mediated currents (20.3 ± 1.5 pA, n=32 spines, one-way ANOVA (F=10.3, p<0.0001), Tukey’s multiple comparison test p>0.05) or NMDAR currents (23.0 ± 2.5 pA, n=25 spines, one-way ANOVA (F=5.01, p=0.0008), Tukey’s multiple comparison test p>0.05) but reduced NMDAR-dependent ΔCa2+ (to 0.39 ± 0.01 ΔG/G_sat_, one-way ANOVA (F=12.11, p<0.0001), Tukey’s multiple comparison test p<0.001, Figures 3B, 4).

**Figure 3.**
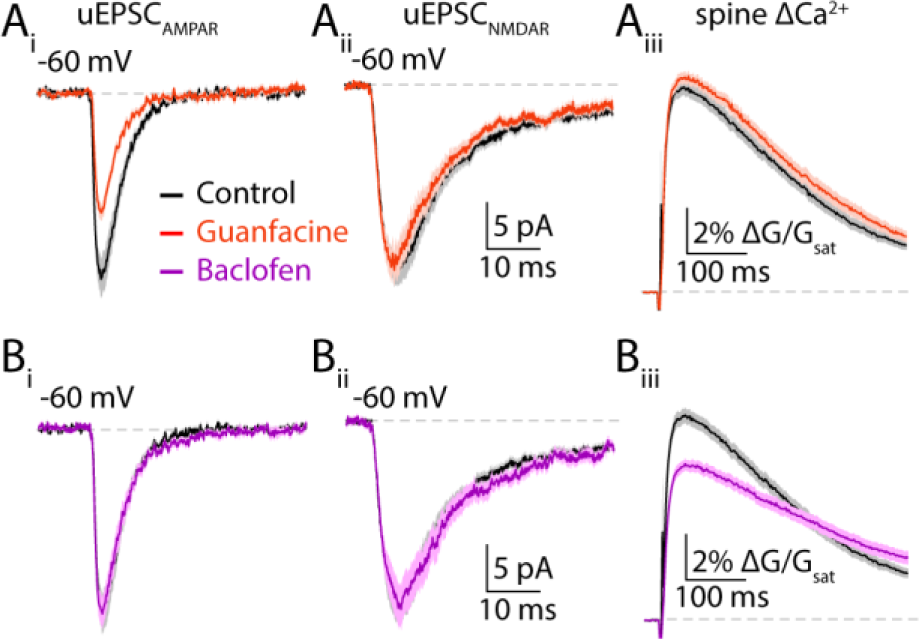
Differential control of excitatory transmission by α2- and GABAb receptors. (Ai) Mean AMPAR-mediated uEPSCs in control (black) and guanfacine (orange) ± SEM (shaded areas). (Aii) 2PLU-evoked NMDAR currents and (Aiii) Ca2+ transients in control (black) and guanfacine (orange), mean (solid lines) ± SEM (shaded areas). (Bi) Mean AMPAR-mediated uEPSCs in control (black) and baclofen (magenta) ± SEM (shaded areas). (Bii) NMDAR-mediated uEPSCs and (Biii) Ca2+ transients in control (black) and baclofen (magenta) ± SEM (shaded areas). *: p<0.05, unpaired t-test.

We then performed similar experiments in mice injected four hours prior to slice preparation with ketamine. In contrast to saline-treated animals, AMPAR-mediated currents were significantly reduced by baclofen (to 12.1 ± 1 pA, n=33 spines, one-way ANOVA (F=9.93, p<0.0001), Tukey’s multiple comparison test p<0.001, Figure 4A) while guanfacine significantly reduced both NMDAR currents (to 13.6 ± 1.3 pA, n=37 spines, one-way ANOVA (F=5.0, p=0.0008), Tukey’s multiple comparison test p<0.05, Figure 4B) and ΔCa2+ (to 0.42 ± 0.02 ΔG/G_sat_, one-way ANOVA (F=12.1, p<0.0001), Tukey’s multiple comparison test p<0.001, Figure 4C). We have previously shown that blocking RGS4 function does not alter the effect of guanfacine on AMPARs or the baclofen induced reduction of NMDAR dependent Ca2+ influx. We have also shown that multiple methods of RGS4 inhibition introduced baclofen induced reduction of NMDAR currents (Lur *et al*, 2015). Our current results match these previous observations. Overall, these findings demonstrate that ketamine exposure disrupts neuromodulation of glutamatergic signaling in the absence of direct actions on synaptic potency.

**Figure 4.**
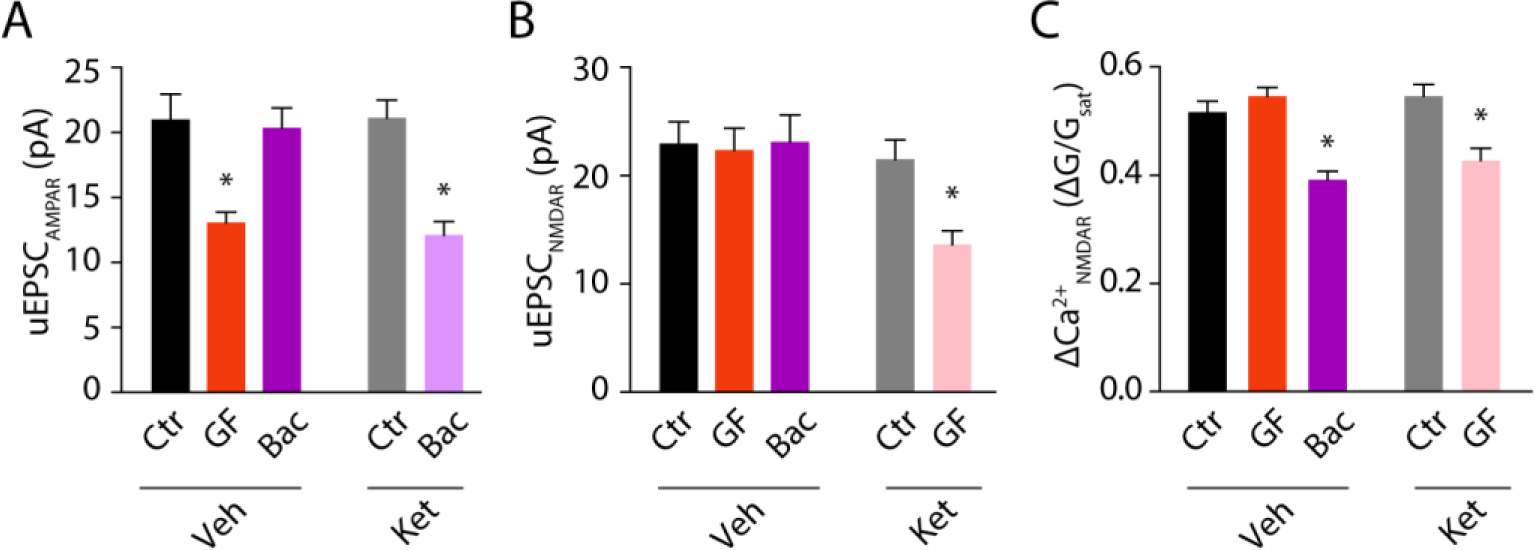
Ketamine treatment eliminates compartmentalized neuromodulation. (A) Bars represent mean amplitude ± SEM of AMPAR currents, (B) NMDARmediated currents and (C) NMDAR-mediated Ca2+ transients in control (gray), in guanfacine (orange) or in baclofen (magenta) from vehicle- versus ketaminetreated mice. *: p<0.05, Tukey’s multiple comparison test.

## Discussion

In our present study, we demonstrate that a single sub-anesthetic dose of ketamine produces significant dysregulation of neuromodulatory control over glutamatergic signaling in the mouse prefrontal cortex. Specifically, following ketamine administration, adrenergic and GABAergic inputs inhibit NMDARs and AMPARs, respectively, a phenomenon that does not occur in untreated animals. Our results suggest this process is mediated by acute reduction in expression levels of the small GTPase RGS4, which we previously showed prevents neuromodulatory cross-talk in dendritic spines (Lur *et al*, 2015). Our findings thus extend our knowledge of targets for ketamine that may contribute to or interact with its antidepressant actions in vivo.

RGS4 is a small GTPase that limits signaling through G protein-coupled receptor pathways by accelerating the hydrolysis of GTP to GDP (Arshavsky and Pugh, 1998; Bansal *et al*, 2007; Zhong *et al*, 2003). Under control conditions, this activity produces microdomains within single dendritic spines that restricts neuromodulatory cascades. For example, adrenergic α2 receptors and GABAB receptors are both coupled to G_i_ proteins that down-regulate cAMP production and PKA activity. Surprisingly, we found that both receptors are present in single spines but are selectively able to negatively modulate AMPARs and NMDARs, respectively (Lur *et al*, 2015). However, when RGS4 is blocked pharmacologically, this microdomain organization breaks down, leading to cross talk between the neuromodulatory signaling cascades (Lur *et al*, 2015; Zhong *et al*, 2003). Remarkably, a single dose of ketamine appears to produce substantial loss of RGS4 expression within a few hours (Stratinaki *et al*, 2013), a finding confirmed in our present study, suggesting this protein is rapidly turned over in cortical neurons. Consistent with our previous results using small molecule inhibitors of RGS4, ketamine-induced down-regulation of RGS4 expression also produces aberrant cross-talk between modulatory signals and glutamate receptors.

Previous studies looking at the effects of ketamine administration 24 hours post-exposure showed increased amplitude for spontaneous EPSCs in layer 5 PNs. This result was linked to an increase in both spine volume and the density of mature spines and attributed to changes in postsynaptic gene expression (Li *et al*, 2010). In contrast, our results indicate that four hours after ketamine administration, basal postsynaptic glutamatergic signaling through either AMPARs or NMDARs is unaltered. This difference may be due to a longer time window required for altered synaptic gene expression to manifest. Since spontaneous mEPSC recordings do not distinguish between pre- and post synaptic effects it is also possible that the ketamine induced changes reported by Li et al. are presynaptic. Our 2PLU experiments circumvent this caveat by exclusively stimulating postsynaptic glutamate receptors.

Acute doses of ketamine have been shown to produce rapid synaptic reorganization in the prefrontal cortex that coincides with antidepressant actions in rodent models (Kavalali *et al*, 2012; Li *et al*, 2011). Interestingly, these effects are thought to be mediated through inhibition of NMDARs by ketamine (Li *et al*, 2011). This hypothesis is supported by evidence that other NMDAR blockers can produce similar synaptic and behavioral effects (Autry *et al*, 2011; Li *et al*, 2011). Loss of NMDAR signaling may activate both mTOR and BDNF signaling pathways that may provide molecular mechanisms for synaptic changes following ketamine (Autry *et al*, 2011; Jourdi *et al*, 2009; Lepack *et al*, 2014; Li *et al*, 2010; Liu *et al*, 2012; Takei *et al*, 2004). Our findings suggest the intriguing possibility that ketamine may also suppress NMDAR activity by broadening the consequences of adrenergic signaling. That is, following ketamine, α2 receptors may further suppress these glutamate receptors. Thus, activation of adrenergic signaling could be an adjunct approach to boost the effects of ketamine. Indeed, guanfacine alone exhibits antidepressant activity in human studies (Mineur *et al*, 2015; Ramos and Arnsten, 2007).

In conclusion, our current findings provide novel evidence that acute ketamine can influence glutamatergic transmission in the mouse prefrontal cortex via down-regulation of RGS4 expression and dysregulation of neuromodulatory signaling. These results expand our view of the downstream actions of ketamine and suggest that inhibition of NMDARs by α2 adrenergic signaling may provide benefit in models of depression when delivered alongside sub-anesthetic doses of ketamine.

## Acknowledgements

The authors wish to thank Dr. Jessica A. Cardin and members of the Higley laboratory for helpful discussions during the preparation of this manuscript.

